# NAb-seq: an accurate, rapid and cost-effective method for antibody long-read sequencing in hybridoma cell lines and single B cells

**DOI:** 10.1101/2022.03.25.485728

**Authors:** Hema Preethi Subas Satish, Kathleen Zeglinski, Rachel T. Uren, Stephen L. Nutt, Matthew E. Ritchie, Quentin Gouil, Ruth M. Kluck

**Author notes:** joint first authors (Hema Preethi Subas Satish, Kathleen Zeglinski). joint last authors (Quentin Gouil, Ruth M. Kluck). Contact: Ruth M. Kluck, Blood Cells and Blood Cancer Division, The Walter and Eliza Hall Institute of Medical Research, Melbourne, Australia.

## Abstract

Despite their common use in research, monoclonal antibodies are currently not systematically sequenced. This can lead to issues with reproducibility and the occasional loss of antibodies with loss of cell lines. Hybridoma cell lines have been the primary means of generating monoclonal antibodies from immunized animals including mice, rats, rabbits and alpacas. Excluding therapeutic antibodies, few hybridoma-derived antibody sequences are known. Sanger sequencing has been “the gold standard” for antibody gene sequencing but relies on the availability of species-specific degenerate primer sets for amplification of light and heavy antibody genes, in addition to lengthy and expensive cDNA preparation. Here we leveraged recent improvements in long-read Oxford Nanopore Technologies (ONT) sequencing to develop NAb-seq: a three-day, species-independent, and cost-effective workflow to characterize paired full-length immunoglobulin light and heavy chain genes from hybridoma cell lines. When compared to Sanger sequencing of two hybridoma cell lines, long-read ONT sequencing was highly accurate, reliable, and amenable to high throughput. We further show that the method is applicable to single cells, allowing efficient antibody discovery in rare populations such as memory B cells. In summary, NAb-seq promises to accelerate identification and validation of hybridoma antibodies as well as antibodies from single B cells used in research, diagnostics and therapeutics.

## INTRODUCTION

Antibodies are essential tools for research as well as for diagnostic and therapeutic applications as they can bind specific targets. However, many research antibodies are poorly characterized, and around half lack specificity for their reported targets ^1, 2^. Poor quality antibodies waste resources and confound scientific studies such that they cannot be reproduced by others ^3-5^. There are very few instances where hybridoma antibody genes have been sequenced, due in part to the added costs and complexity of antibody sequencing.

To improve the reliability of antibodies, the Antibody Society ^6^ and leading scientists ^4, 7^ have recommended collaboration and funding to define antibodies by their DNA sequence. Sequencing provides a basis on which to validate antibody specificity and sensitivity across all relevant applications. It also allows the generation of well-defined recombinant antibodies by incorporating the genes into plasmid DNA followed by expression in mammalian or bacterial cells ^8^.

Conventional sequencing of antibody genes from hybridoma cell lines involves PCR amplification of antibody variable regions (V_H_ and V_L_) followed by low-throughput and partial-length Sanger sequencing. Primer sets for antibody gene amplification are readily available for mouse and human loci but are less available for other species thereby limiting broader application of this sequencing approach. In addition, Sanger sequencing by commercial providers costs around US$800 (2 weeks) for the variable regions (V_H_ and V_L_) and US$2,000 (4 weeks) for the full-length antibody (variable and constant regions). Even when performing Sanger sequencing in-house, there is still an estimated turnaround time of five days and cost of US$70-120 per antibody ^9^. Alternatively, Illumina sequencing has been incorporated into antibody discovery platforms for high-throughput short-read sequencing of antibody heavy and light chains ^10-13^. However, these protocols also rely on species-specific primers and generate partial-length reads (up to 600 bp) which require assembly, while the high throughput is not well-suited to small-scale monoclonal antibody sequencing for a limited number of cell lines.

Long-read sequencing allows for the full-length sequencing of antibodies, however a relatively high error rate compared with short-read sequencing approaches has limited its application in antibody sequencing thus far ^14^. PacBio long-read sequencing has been applied to the sequencing of single B cells ^15^ and phage display libraries ^16, 17^, but its high cost makes it difficult to implement routinely. By contrast, Oxford Nanopore Technologies (ONT) sequencing has a much lower capital cost and flexible throughput, with raw-read accuracy having improved in recent years to >95%. The remaining errors in ONT, notably systematic homopolymer errors (insertions/deletions in tracts of >5 identical bases) can lead to frameshifts, complicating translation of nanopore-derived sequences into accurate antibody protein sequences ^18^. Notably, improvements in sequencing accuracy and correction techniques ^19^ have enabled the study of antibody fragments ^20^ and B cells ^21-23^, although these studies employ complex and costly error correction methods ^22^, are specific to cancerous cells ^21^ or are too high throughput to be realistically used for hybridoma sequencing ^24^.

To bring about the transition to sequence-defined recombinant antibodies, we developed NAb-seq (Nanopore Antibody sequencing), a simplified experimental and computational workflow based on ONT sequencing to obtain full-length antibody sequences from two rat hybridoma cell lines and compared to outsourced Sanger sequencing results. One million full-length cDNA reads were generated from multiplexed hybridomas on an ONT Flongle flow cell and assembled into 100% accurate antibody chains. With feasibility to upscale to 24 hybridomas per flow cell, consumables are estimated at approximately US$30 per antibody within a 3-day turnaround (1 day library preparation, 1 day sequencing, 1 day analysis). We further show that NAb-seq can be applied to the antibody sequencing of single B cells. NAb-seq is thus a valuable tool for recombinant antibody production, humanization of therapeutic antibodies, and protection of intellectual property.

## RESULTS

### Long-read sequencing of antibody genes from Rattus norvegicus hybridoma cells

Antibody sequencing requires high accuracy due to the crucial role of somatic mutation of variable regions in antibody specificity and affinity for the target. Recent improvements in the accuracy of ONT long-read data prompted us to test its efficacy in rapid sequencing of hybridoma antibody genes. We tested two hybridoma cell lines that had been developed in-house and whose antibody genes had been Sanger sequenced by commercial sources.

The workflow for NAb-seq is outlined in Figure 1. cDNA libraries A and B were prepared from total RNA of two hybridoma cell lines using the ONT PCR-cDNA barcoding kit. Analysis of the libraries in a DNA TapeStation (Figure 2a) showed bands at 1600 and 900 kb, the expected size for full-length antibody heavy and light chain. The two cDNA libraries were then pooled for parallel long-read sequencing using the ONT Flongle flow cell, which generated ∼1 million raw reads in 24 hours. The sequence data were basecalled in super-high accuracy mode and aligned to the reference *Rattus norvegicus* antibody gene sequences obtained from the international ImMunoGeneTics information system^®^ (IMGT, http://www.imgt.org/ligmdb/) ^25^. The number of pass (Q score >10) reads were ∼ 300,000 and 200,000 for library A and B (Figure 2b), with read length distribution shown in Figure 2c. Thus, even without enrichment, 2.04% of pass reads in library A and 3.51% of pass reads in library B corresponded to antibody transcripts (Table 1), providing more than enough antibody reads for consensus sequence generation. Empirically, as few as 5 reads could generate a 100% accurate consensus for a heavy chain. This means that there is scope to multiplex up to 24 hybridomas per Flongle flow cell (using the recent SQK-PCB111.24 nanopore library preparation kit) and still generate ∼42,000 reads per hybridoma with 2-3% (∼800-1200) of those being antibody transcripts.

**Table 1.**
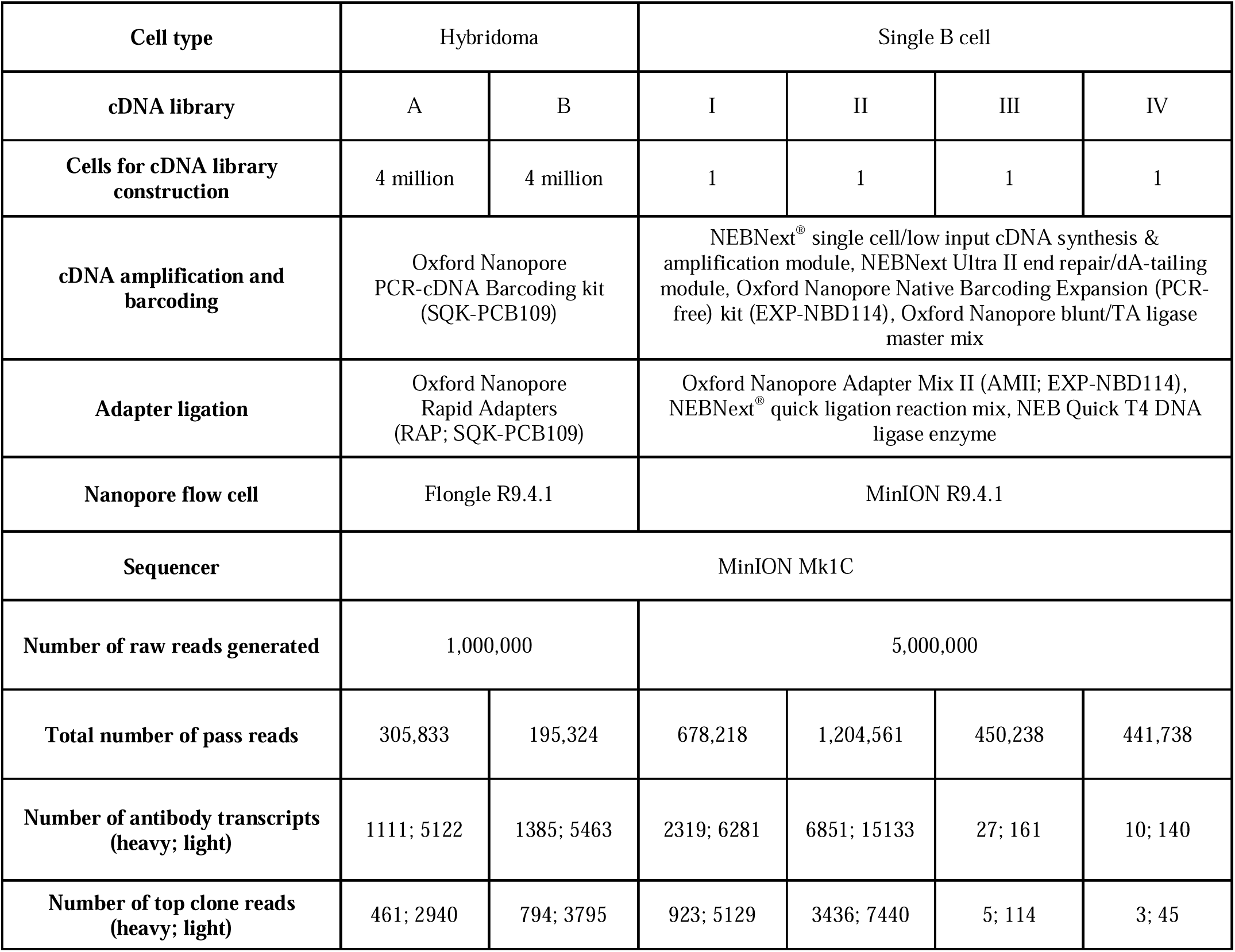
Summary of antibody gene sequencing using NAb-seq

| Cell type | Hybridoma |  | Single B cell |  |  |  |
| --- | --- | --- | --- | --- | --- | --- |
| cDNA library | A | B | I | II | III | IV |
| Cells for cDNA library construction | 4 million | 4 million | 1 | 1 | 1 | 1 |
| cDNA amplification and barcoding | Oxford Nanopore PCR-cDNA Barcoding kit (SQK-PCB109) |  | NEBNext® single cell/low input cDNA synthesis & amplification module, NEBNext Ultra II end repair/dA-tailing module, Oxford Nanopore Native Barcoding Expansion (PCR-free) kit (EXP-NBD114), Oxford Nanopore blunt/TA ligase master mix |  |  |  |
| Adapter ligation | Oxford Nanopore Rapid Adapters (RAP; SQK-PCB109) |  | Oxford Nanopore Adapter Mix II (AMII; EXP-NBD114), NEBNext® quick ligation reaction mix, NEB Quick T4 DNA ligase enzyme |  |  |  |
| Nanopore flow cell | Flongle R9.4.1 |  | MinION R9.4.1 |  |  |  |
| Sequencer | MinION Mk1C |  |  |  |  |  |
| Number of raw reads generated | 1,000,000 |  | 5,000,000 |  |  |  |
| Total number of pass reads | 305,833 | 195,324 | 678,218 | 1,204,561 | 450,238 | 441,738 |
| Number of antibody transcripts (heavy; light) | 1111; 5122 | 1385; 5463 | 2319; 6281 | 6851; 15133 | 27; 161 | 10; 140 |
| Number of top clone reads (heavy; light) | 461; 2940 | 794; 3795 | 923; 5129 | 3436; 7440 | 5; 114 | 3; 45 |

**Figure 1.**
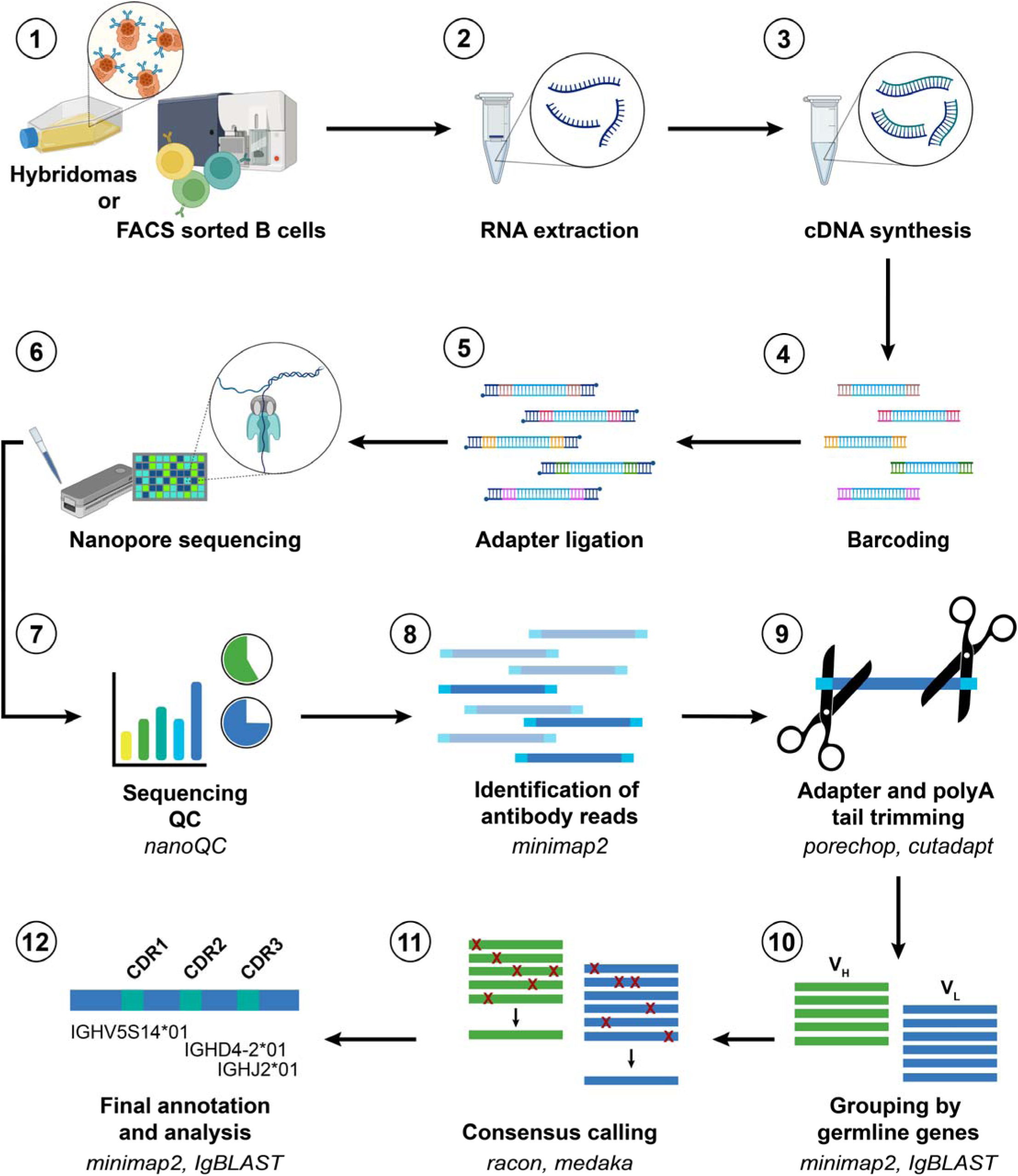
NAb-seq workflow for parallel sequencing of full-length antibody heavy and light chain sequences from hybridoma cell lines and single B cells.

**Figure 2.**
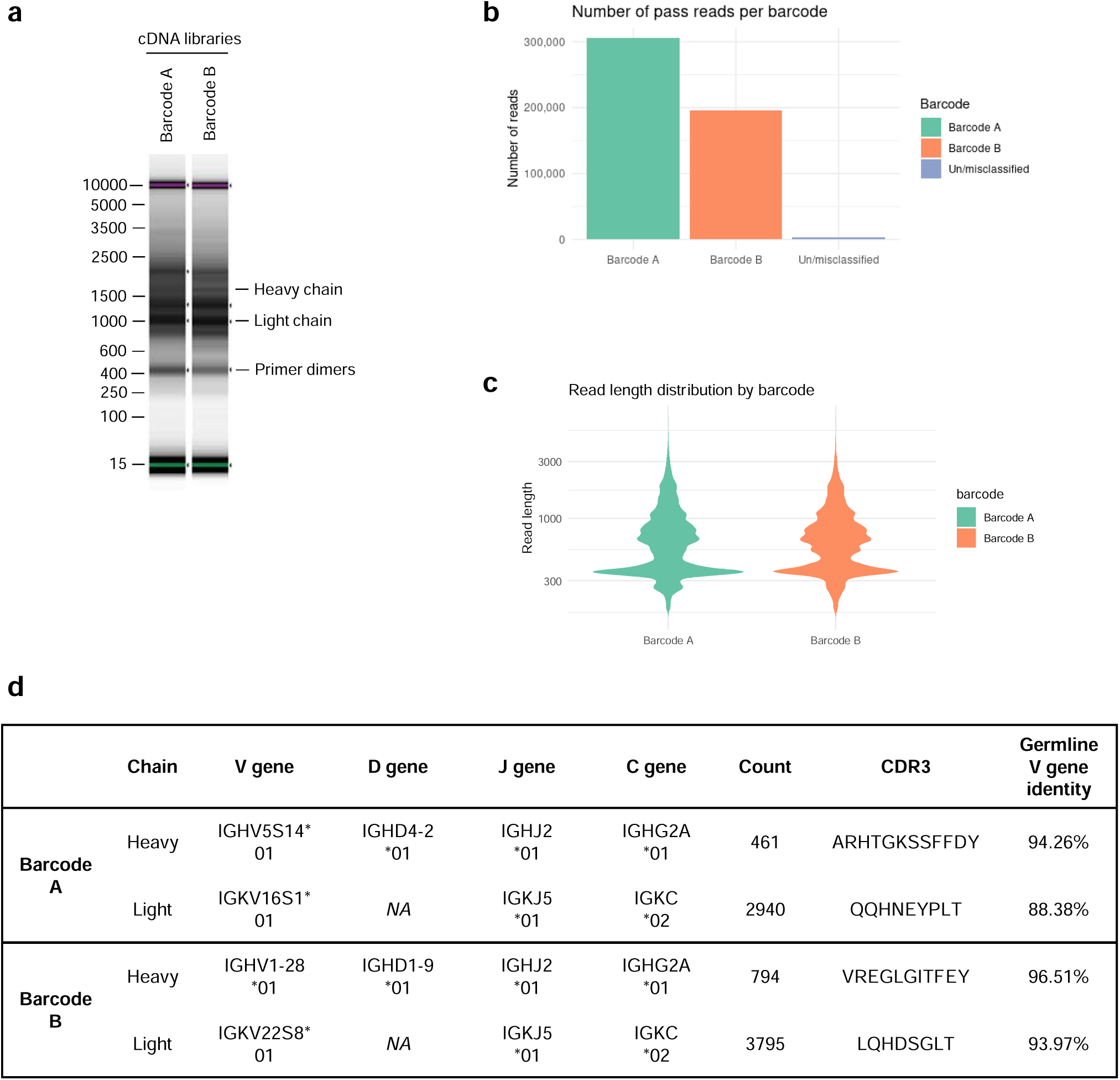
NAb-seq of two hybridoma cell lines revealed antibody sequences. (a) cDNA library size and amplification. Total RNA from two hybridoma cell lines were extracted and converted to cDNA followed by amplification and barcoding, generating whole transcriptome cDNA libraries A and B. (b) Basecalling in Guppy’s super-high accuracy mode yielded approximately 0.5 million total pass reads. (c) Read length of pass reads varied from ∼400 bp to ∼3000 bp. (d) Sequence analysis of cDNA libraries A and B reveals V(D)J recombination, C gene usage and complementarity determining region 3 (CDR3) amino acid sequence. Consensus calling of antibody transcripts revealed IgG isotype.

To generate accurate full-length antibody sequences, reads trimmed of their polyA tails were aligned against germline antibody sequences using three tools: IgBLAST^26^, IMGT/V-QUEST^25^ and minimap2^27^. Only antibody transcripts with identical V(D)J and C genes were grouped together for consensus calling to avoid generating a chimeric consensus. The consensus sequences with the most abundant V(D)J and C gene combinations were taken as the heavy and light chain sequences for each cell. Dominant gene combinations were typically 5-30x more abundant than the second most common combinations (heavy chains were typically 5-10x, light chains were typically 10-30x). The heavy chain isotype of both hybridomas was IgG2A (Figure 2d).

The heavy and light chain sequences from libraries A and B aligned with 100% accuracy to the Sanger-generated sequences of 7D10 and 3C11, respectively (Figure 3 a,b and Supplementary Figures S1-3). Further analysis using BLAST^28^ showed that additional sequences at the 5’ and 3’ ends of the heavy and light chains matched with the 5’ and 3’ UTR sequences of a *Rattus norvegicus* antibody.

**Figure 3.**
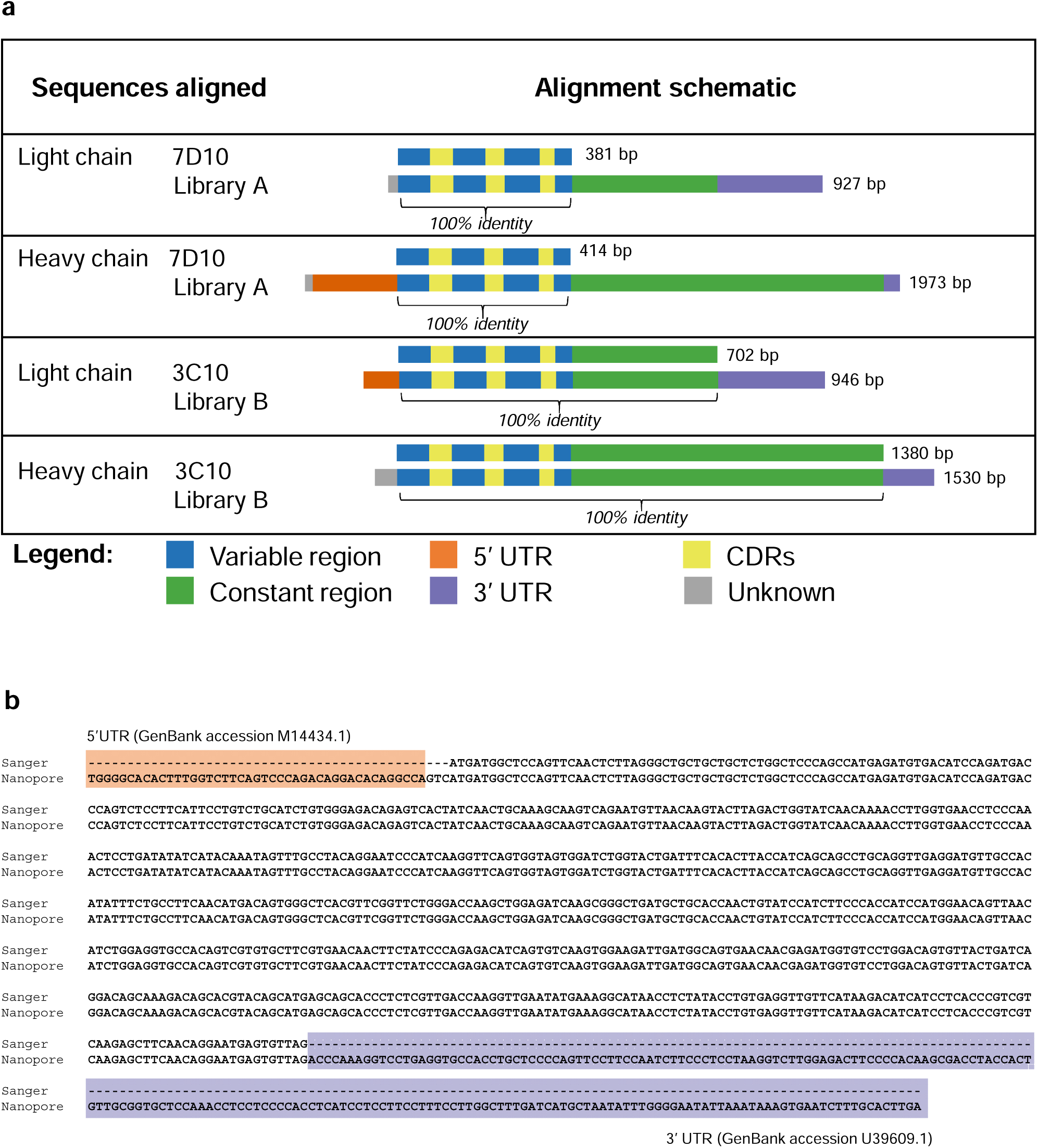
Antibody sequences from libraries A and B align with 100% accuracy to the 7D10 and 3C10 sequences. (a) Schematic of alignment of 7D10 and 3C10 antibody chains, as derived from Sanger (top row) and Nanopore (bottom row) sequencing methods. Additional bases present in the Nanopore sequence have been annotated with BLAST as indicated. Short sequences at the beginning of reads (grey) were sometimes unable to be annotated with BLAST, nor did they match the primer sequences used during library preparation. (b) Sequence alignment of the light chain sequence of 3C10 as derived from Sanger (top row) and Nanopore (bottom row) sequencing methods. Additional bases present in the Nanopore sequence have been annotated with BLAST. Regions highlighted as in Figure 3a. See Supplementary Figures 1-3 for other chains of 7D10 and 3C10.

### Long-read sequencing of antibody genes from Rattus norvegicus single B cells

Following the successful application of NAb-seq to bulk hybridoma cell line samples, we assessed if this approach could also recover antibody genes from single primary cells that produce antibodies, such as purified rat B cells. Splenocytes from rats immunized with BAX peptide were harvested, enriched, and sorted to isolate B cells with antibodies specific for the corresponding region in BAX (for further details see Methods).

Four individual B cells were sorted, lysed, and cDNA nanopore libraries (I-IV) were prepared from total RNA using the NEBNext^®^ Single cell/Low Input kit followed by native barcoding. Bands at the expected size for full-length antibody heavy and light chain genes were not evident after analysis on a DNA TapeStation (Figure 4a), suggesting low expression of antibody genes. The cDNA libraries were sequenced using a MinION flow cell to provide increased sequencing depth over the Flongle, as we could not predict the expression levels of antibody genes nor the efficiency of the cDNA preparation. Accordingly, ∼5 million pass reads (Q score >10) were generated during a 72 hour run (Figure 4b-c). Basecalling was again performed in super-high accuracy mode.

**Figure 4.**
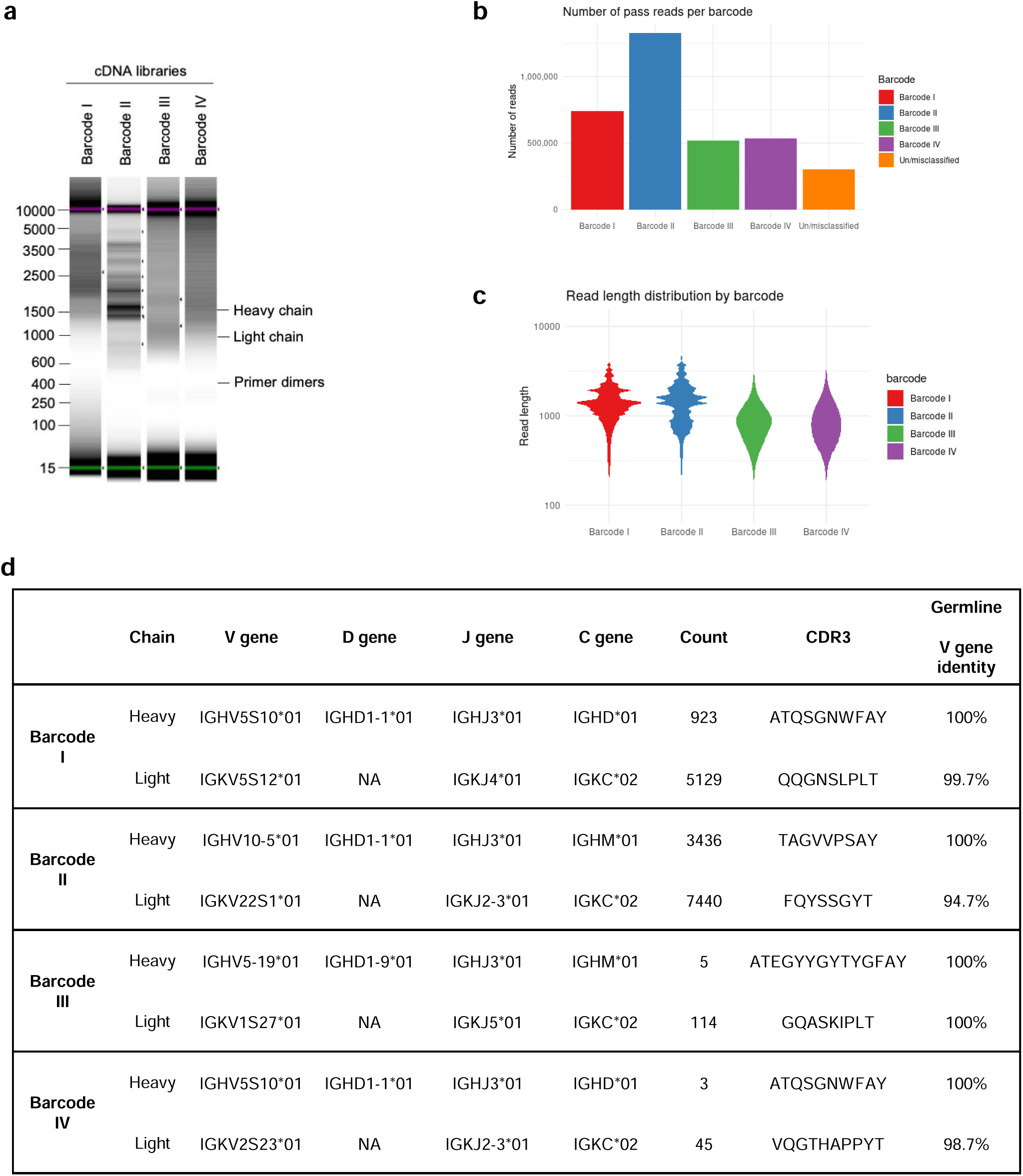
NAb-seq of four single B cells revealed antibody sequences. (a) cDNA library size and amplification. Total RNA from four sorted single B cells were extracted and converted to whole transcriptome cDNA libraries I-IV. (b) Basecalling in Guppy’s super-high accuracy mode yielded approximately 5 million total pass reads. (c) Read length of pass reads varied from ∼400 bp to ∼5000 bp. (d) Sequence analysis of cDNA libraries I to IV reveals V(D)J recombination, C gene usage and complementarity determining region 3 (CDR3) amino acid sequence. Consensus calling of antibody transcripts revealed IgM and IgD isotypes.

Alignment of the pass reads to the reference *Rattus norvegicus* antibody gene sequences deposited in the international ImMunoGeneTics information system^®^ (IMGT ^25^) revealed that 1.27% of pass reads in library I and 1.82% of pass reads in library II corresponded to antibody transcripts (Table 1). However, a low percentage of pass reads in libraries III and IV corresponded to antibody transcripts (0.04% and 0.03%, respectively). The low number of antibody transcripts in cDNA libraries III and IV is a consequence of less efficient PCR amplification resulting in a high proportion of artefacts, with few reads overall aligning to the rat transcriptome (7.28% for library III and 3.26% for library IV vs ∼80% for libraries I and II, Table 1). Nonetheless, more than 100 antibody reads were obtained for each library, and consensus sequences could be generated for all four libraries. Upon alignment to germline sequences using IMGT/V-QUEST^25^, the isotype of the heavy chains was identified to be either IgM or IgD, rather than IgG (Figure 4d). This indicated that all 4 cells were likely to be naïve, rather than mature switched memory B cells. In addition, consensus calling revealed near 100% identity with germline sequences (Figure 4d), indicative of antibodies expressed on naïve B cells. The small degree of sequence mismatch observed in the light chain V regions is likely due to genetic variation between the rat strain used for this study (Wistar) and the rat reference genome generated in the BN/SsNHsd strain (Figure 4d). Regardless, these data show that, in addition to the characterization of hybridoma cell lines, the NAb-seq workflow could identify antibody sequence and isotype with near 100% accuracy from individual B cells and quickly determine that the sorting procedure for antibodies specific to BAX had failed.

## DISCUSSION

The widespread adoption of high-quality recombinant antibodies is essential for improving biological research quality and reproducibility and requires an accurate and cost-effective means of sequencing antibody genes from pre-existing and future hybridomas. Here we have developed a new long-read sequencing workflow, NAb-seq, to provide full-length sequences of antibody heavy and light chains from hybridoma cell lines, as well the sequence of antibody genes from single naïve rat B cells. High-quality, in-frame antibody consensus sequences were generated by basecalling raw reads using guppy’s super high accuracy model, followed by a two-step consensus-calling process using Racon^29^ and Medaka (https://github.com/nanoporetech/medaka) for the removal of sequencing errors, including those associated with homopolymers. This was achieved with as few as five copies, an improvement on previous studies which indicated up to 25 may be required ^19^. Thus, NAb-seq generated antibody gene sequences with perfect accuracy (100%).

By sequencing the whole transcriptome, NAb-seq offers a simple protocol that obviates the need for species-specific customized primers. NAb-seq could thus be used for the determination of antibody sequences from any species. This simple and universal protocol is a major advantage over all targeted sequencing techniques (Sanger, Illumina, PacBio/ONT; ^9, 10, 17, 20^). It is also much simpler, cheaper and less prone to chimerism than whole-transcriptome concatemer nanopore sequencing ^22^.

The trade-off for the whole-transcriptome sequencing strategy is that most of the reads are not antibody transcripts and are therefore discarded. Still, we found that about 2% of the reads sequenced from hybridoma cell lines or single B cells correspond to antibody transcripts. Combined with the capacity of a nanopore Flongle flow cell to generate in excess of a million reads, this allows for multiplexing dozens of samples in a single run. The use of sample barcodes in NAb-seq also allows for the pairing of heavy and light chain sequences in each sample. The achieved throughput is far superior to Sanger sequencing, while the ability to tune sequencing throughput (using different size nanopore flow cells, washing and reusing flow cells) makes it more flexible than Illumina or PacBio sequencing.

NAb-seq also benefits from a streamlined analysis workflow. By contrast to Illumina sequencing, long reads span the entire antibody transcript, which greatly simplifies the bioinformatic analysis workflow as there is no need for assembly ^30^. With improvements in basecalling accuracy and robust error correction ^19^, NAb-seq improves on previous efforts to sequence antibodies with nanopore by using a simple consensus calling approach to error correction that does not require additional library preparation steps or computationally intensive bioinformatic steps. Lowden and Henry ^20^ found that, without error correction steps, 75-80% of antibody fragment reads were unable to have their CDR3 identified. Error correction methods that rely on the sequencing of concatemers from rolling circle amplification can successfully reconstruct BCR sequences from single B cells ^22^, however they require more time and computational power than NAb-seq. Similarly, the assembly-based approach employed to characterize cancerous B cells ^21^, though effective, is slower and more computationally intensive than the simple consensus strategy implemented by NAb-seq.

In summary, NAb-seq is ideally suited for quickly and cheaply generating high-accuracy antibody sequences from between 1 and 24 hybridoma cell lines or single B cells. The affordability of the minION nanopore sequencing instrument (US$1000), low cost per run, universal protocol, flexible throughput and quick turnaround enable NAb-seq to be performed in-house and easily integrated into existing workflows. For much larger sample numbers (e.g. hundreds or thousands of hybridomas or single B cells using 10X single-cell approaches), targeted methods like RAGE-Seq which sequence only heavy/light chain amplicons become more efficient ^24^. Still, the computational error correction (consensus) approach implemented in NAb-seq could be applied to targeted methods to generate high accuracy consensus sequences.

In summary, NAb-seq could help to deliver well-defined antibody products for research, diagnostic and therapeutic applications. In addition, it could facilitate the supply of all antibodies as recombinant proteins to improve reproducibility, an approach some manufacturers have started to implement ^7^.

## MATERIALS AND METHODS

### Hybridoma cell culture and total RNA extraction

Hybridomas expressing the rat monoclonal antibody clones 7D10 and 3C11 (WEHI Antibody Facility) were cultured in Hybridoma serum-free medium (12045076; Gibco) supplemented with 10% foetal bovine serum (FBS; F9423; Sigma) and 100 U/ml interleukin-6 at 37 °C in 8% humidified CO_2_. Hybridomas were harvested and spun down at 300 x g for 5 min. Pelleted cells were resuspended in sterile 1x Dulbecco’s phosphate-buffered saline (DPBS; 14190144; Gibco). Total RNA was extracted from approx. 4 × 10^6^ cells using the Qiagen RNeasy mini kit (74104; Qiagen) as per the manufacturer’s protocol and quantified using DeNovix DS-11 series spectrophotometer.

### cDNA synthesis and library construction from hybridomas

Whole transcriptome cDNA libraries were constructed from extracted mRNA using a PCR-cDNA barcoding kit (SQK-PCB109; Oxford Nanopore Technologies). Briefly, 50 ng total RNA from each hybridoma was taken in RNase-free PCR tubes containing Maxima H Minus reverse transcriptase enzyme (EP0751; ThermoFisher Scientific), VN primers and strand-switching primers and incubated at 42 °C for 90 min (one cycle) followed by enzyme inactivation at 85 °C for 1 min (one cycle) in a thermocycler. This was followed by full-length cDNA amplification and barcoding via 12 PCR cycles using barcoded primers and 2x LongAmp Taq master mix (M0287; New England Biolabs). PCR cycling conditions were set as: initial denaturation at 95 °C for 30 s (1 cycle), denaturation at 95 °C for 15 s (12 cycles), annealing at 62 °C for 15 s (12 cycles), extension at 65 °C for 4 min 10 s (12 cycles) and final extension at 65 °C for 6 min (1 cycle). To selectively amplify cDNAs of length up to ∼5 kb the extension time was adjusted to 4 min 10 s (50 s per kb). PCR reactions with the same barcode were pooled in 1.5 ml Eppendorf^®^ DNA LoBind tubes (EP0030108051; Merck) and cleaned up using 0.8x equivalent Agencourt^®^ AMPure^®^ XP beads (A63880; Beckman Coulter). The cleaned-up cDNA libraries were quantified using Qubit dsDNA HS kit (Q32851; ThermoFisher Scientific). Size distribution of the amplified cDNAs was analysed using High Sensitivity D5000 ScreenTape (50675592; Agilent Technologies) in a 4200 TapeStation (Agilent).

### Long-read sequencing of hybridoma cDNA libraries

Long-read sequencing also used the PCR-cDNA barcoding kit (SQK-PCB109; Oxford Nanopore Technologies). Before loading ∼50 fmol of cDNA libraries, a Flongle flow cell R9.4.1 (FLO-FLG001; Oxford Nanopore Technologies) in a minION Mk1C device was primed with pre-mixed flush buffer and flush tether (EXP-FLP002; Oxford Nanopore Technologies). For ligation of cDNA libraries with adaptors, the libraries (100 fmol of each in a volume of 11 µl) were pooled in a 1.5 ml Eppendorf^®^ DNA LoBind tube, followed by addition of rapid adapters and incubation at room temperature for 5 min. The sequencing mix was prepared in a fresh 1.5 ml Eppendorf^®^ DNA LoBind with sequencing buffer, loading beads and the pooled barcoded cDNA libraries. After quality checking the flow cell, the sequencing mix was added to the flow cell and run for 24 hours. Upon completion, guppy (v5.0.11) was used to perform super high accuracy basecalling and demultiplexing of the data.

### Single rat B cell sort

Single rat B cells were sourced from a separate project designed to generate antibodies to a specific region in the pro-apoptotic protein BAX. Two Wistar rats had been immunized with KLH-conjugated BAX peptide and splenocytes used to generate hybridomas, with the excess frozen. As the hybridomas had not generated the desired antibodies, we pursued the possibility that B cell cloning could sort rare memory B cells that generated site-specific antibodies to BAX. As rat B cell cloning had not been reported, the mouse B cell cloning procedure ^31^ was modified to sort rat B cells. In addition, long-read sequencing was pursued to avoid the need for rat-specific primers.

Splenocytes were rapidly thawed in a 37 °C water bath, pelleted and resuspended in FACS buffer (1x DPBS supplemented with 2% FBS and 1 mM EDTA) and stored on ice. Following cell counting, the splenocytes were centrifuged at 300 x g for 4 min and resuspended in FACS buffer at 5 × 10^7^ cells/ml. The cells were then enriched for B cells using EasySep^®^ rat B cell isolation kit (19644; STEMCELL Technologies) according to manufacturer’s instructions.

To select for memory B cells, and for those that bind to the BAX peptide, the enriched cells were resuspended at 10-20 million cells per 100 ml and incubated at 4 °C with the following mixture of antibodies: mouse anti-rat CD32 antibody (rat BD Fc block; 2 ml per 100 ml cells; 556970; BD Biosciences); FITC anti-rat IgM (2 ml per 100 ml cells; 408905; BioLegend); PE anti-rat CD3 (2.5 ml per 100 ml cells; 201411; BioLegend); PE anti-rat CD11b/c (1.25 ml per 100 ml cells; 201807; BioLegend); APC anti-rat CD45RA (1.25 ml per 100 ml cells; 202314; BioLegend); avi-BAX peptide conjugated to BV 510 streptavidin (SA/BV510; BioLegend); avi-BAX peptide conjugated to BV 785 streptavidin (SA/BV785; BioLegend). Prior to FACS sorting, cells were washed twice with FACS buffer and resuspended at 8 × 10^6^ cells/ml. Cells that were IgM^-^ CD3^-^ CD11b/c^-^ CD45RA^+^ BAX^+^ were sorted using BD FACSAria III into 1x DPBS in a 96-well semi skirt PCR plate (342000; Interpath Services), sorting a total of 4 single B cells.

### cDNA synthesis and library preparation from single rat B cells

Whole transcriptome cDNA library from the sorted single B cells were prepared using the NEBNext^®^ single cell/low input cDNA synthesis & amplification module (E6421S; New England BioLabs Inc.). NEBNext^®^ cell lysis buffer (10X), murine RNase inhibitor and nuclease-free water provided in the kit were added to wells with single B cells and incubated at room temperature for 5 min to lyse cells. NEBNext^®^ single cell RT primer mix was added to each lysed cell and mixed well by pipetting. The plate was incubated in a thermocycler at 70 °C for 5 min for primer annealing and NEBNext^®^ single cell RT buffer, NEBNext^®^ template switching oligo, NEBNext^®^ single cell RT enzyme mix, and nuclease-free water were added for reverse transcription and template switching. Reverse transcription and template switching reactions were incubated 42 °C for 90 min followed by enzyme inactivation at 70 °C for 10 min in a thermocycler. The resulting cDNAs were amplified via 21 PCR cycles using NEBNext^®^ single cell cDNA PCR master mix and NEBNext^®^ single cell cDNA PCR primer mix. The PCR cycling conditions were set up as: initial denaturation at 98 °C for 45 s (1 cycle), denaturation at 98 °C for 10 s (21 cycles), annealing at 62 °C for 15 s (21 cycles), extension at 72 °C for 6 min (21 cycles) and final extension at 72 °C for 5 min (1 cycle). The single cell RT primer mix, single cell RT buffer, template switching oligo, single cell RT enzyme mix, single cell cDNA PCR master mix and single cell cDNA PCR primers were provided in the kit (E6421S; New England BioLabs Inc.). To selectively amplify long transcripts of cDNA, the extension time was adjusted to 6 min. The libraries were cleaned up using 0.8x equivalent Agencourt^®^ AMPure^®^ XP beads. The cleaned-up cDNA libraries were quantified using Qubit dsDNA HS kit. To obtain sufficient cDNA for barcoding, a second round of PCR amplification was performed with 15 cycles and extension time of 7 min. The cDNA libraries were taken in fresh 1.5 ml Eppendorf^®^ DNA LoBind tubes and end-prepped for barcoding using NEBNext^®^ Ultra II end repair/dA-tailing module (E7546S; New England BioLabs Inc.). The end-prep reactions were incubated at 20 °C for 15 min followed by enzyme inactivation at 65 °C for 15 min. The end-prepped cDNA libraries were cleaned up using 1x equivalent Agencourt^®^ AMPure^®^ XP beads. The cleaned-up end-prepped cDNA libraries were quantified using Qubit dsDNA HS kit. The end-prepped cDNA libraries were taken in individual fresh 1.5 ml Eppendorf^®^ DNA LoBind tubes and barcoded using native barcode primers (EXP-NBD114; Oxford Nanopore Technologies) and blunt/TA ligase master mix (M0367S; Oxford Nanopore Technologies). The barcoding reactions were incubated at room temperature for 30 min and cleaned up using 1x equivalent Agencourt^®^ AMPure^®^ XP beads. The barcoded libraries were quantified and pooled (∼100-200 fmol final). Adapters were ligated to the pooled library using adapter mix II (AMII; EXP-NBD114; Oxford Nanopore Technologies), NEBNext^®^ quick ligation reaction mix (E6056S; New England BioLabs Inc.) and Quick T4 DNA ligase (E6056S; New England BioLabs Inc.), by incubation at room temperature for 10 min. The adapter ligated libraries were cleaned up using 1x equivalent Agencourt^®^ AMPure^®^ XP beads and quantified as for the hybridoma libraries.

### Long-read sequencing of single B cell cDNA libraries

Long-read sequencing used the Ligation sequencing kit (SQK-LSK109; Oxford Nanopore Technologies). Before loading of the cDNA libraries onto a MinION R9.4.1 flow cell (FLO-MIN106D; Oxford Nanopore Technologies) in a MinION Mk1C device, the flow cell was primed as for the Flongle flow cell above. The sequencing mix was prepared in a fresh 1.5 ml Eppendorf^®^ DNA LoBind with sequencing buffer loading beads (provided in the SQK-LSK109 kit) and the pooled libraries. After quality checking the flow cell, the sequencing mix was added to the flow cell and run for 72 hours. Upon completion, guppy (v5.0.11) was used to perform super high accuracy basecalling and demultiplexing of the data.

### Bioinformatic analysis

Quality control checks of read counts and length were performed using NanoComp (v1.12.0 ^32^) and polyA tails were trimmed using cutadapt (v3.4 ^33^). All pass (Q score > 10) reads were aligned to reference *Rattus norvegicus* antibody gene sequences retrieved from the international ImMunoGeneTics information system^®^ (IMGT ^25^) using minimap2 (v2.17-r941 ^27^) with the map-ont preset. Any read with an alignment to a germline antibody gene (variable or constant) was considered an antibody transcript and taken forward for annotation. Antibody transcripts were annotated using IgBLAST (v1.17.1 ^26^) with default parameters, and the results written out in AIRR format for processing in R (v4.0.5 ^34^).

In order to correct sequencing errors, antibody transcripts with identical V(D)J and C genes were grouped, and a separate consensus was called for each of the most abundant groups in each cell. Grouping the reads by their germline genes prevents the consensus from being perturbed by transcripts that do not represent the antibody actually expressed by the cell (e.g. resulting from leaky transcription from the second allele or PCR chimeras). Error-corrected consensus sequences were generated using a two-step process: one round of Racon (v1.4.3, using the parameters -w 5000 -t 4 -u -g -8 -x -6 -m 8 --no-trimming ^29^) followed by one round of Medaka (v1.4.2, using default parameters (https://github.com/nanoporetech/medaka)). Finally, the error corrected heavy and light chain sequences for each cell were re-analysed using IgBLAST (v1.17.1 ^26^), minimap2 (v2.17-r941, to identify constant genes ^27^) and the IMGT/V-QUEST ^25^ webserver in order to determine their properties (e.g. percentage identity to germline genes, CDR sequences). All code is available on GitHub (https://github.com/kzeglinski/nab-seq).

## Acknowledgments

We thank Sarah MacRaild, Stephen Wilcox and the WEHI Genomics Platform for help with ONT sequencing, and Paul Masendycz, Kaye Wycherley and the WEHI Antibody Facility for providing hybridoma cell lines.

## Author contributions

HPSS, RU, KZ, QG and RK conceived and designed the study. HPSS and RU assembled the team. HPSS performed the experiments and KZ the analysis. SN consulted the possible applications. RU, SN and MR advised throughout the study. HPSS, KZ, and RK wrote the manuscript, RU, SN, MR, and QG edited and contributed to the final draft. All authors reviewed and approved the final draft.

## Supplementary material

Fig S1

Fig S2

Fig S3

## Data availability statement

All data generated in this study (FASTQ and FAST5) is available from the European Nucleotide Archive (ENA) under accession number PRJEB51442.

## Abbreviations

APC: allophycocyanin
BAX: Bcl-2-associated X protein
BLAST: basic local alignment search tool
BV: brilliant violet
DPBS: Dulbecco’s phosphate-buffered saline
FBS: fetal bovine serum
Ig: immunoglobulin
IMGT^®^: international ImMunoGeneTics information system^®^
KLH: keyhole limpet hemocyanin
NAb-seq: nanopore antibody sequencing
ONT: Oxford Nanopore Technologies
PE: phycoerythrin
RAGE-seq: Repertoire and Gene Expression by Sequencing
V_H_: variable heavy chain
V_L_: variable light chain
V-QUEST: V-QUEry and Standardization

