## Supplementary Data for "NAb-seq: an accurate, rapid and cost-effective method for antibody long-read sequencing in hybridoma cell lines and single B cells"

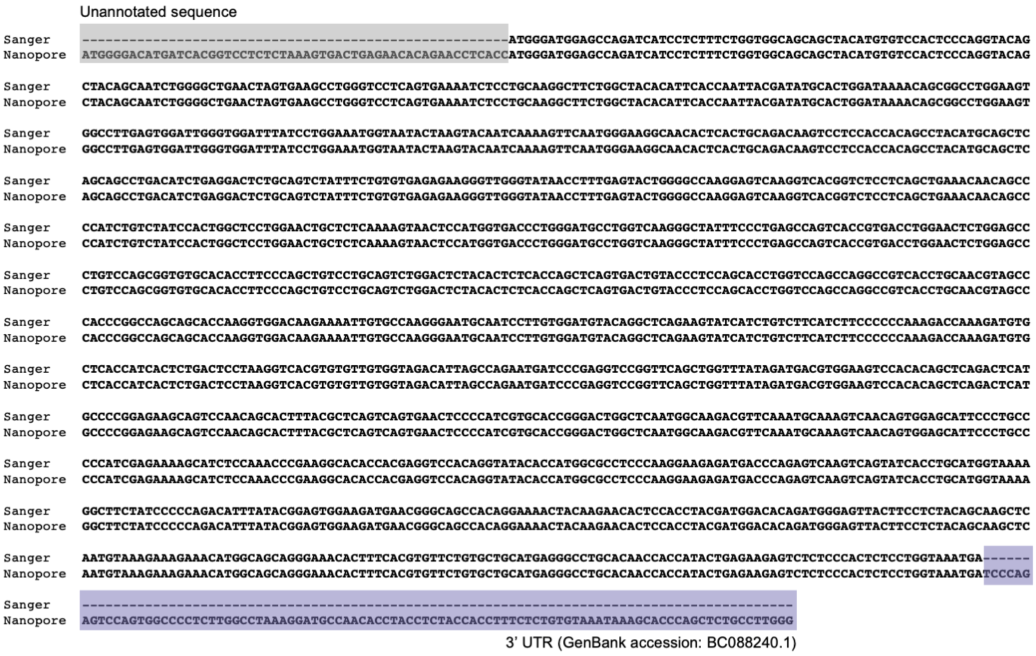


**Supplementary Figure 1.** Alignment of the heavy chain sequence of 3C10, as derived from Sanger (top row) and Nanopore (bottom row) sequencing methods. Additional bases present in the Nanopore sequence have been annotated with BLAST and highlighted as in Figure. 3a. The additional 5' sequence was unable to be annotated with BLAST, nor did it match any of the primer sequences used during library preparation.

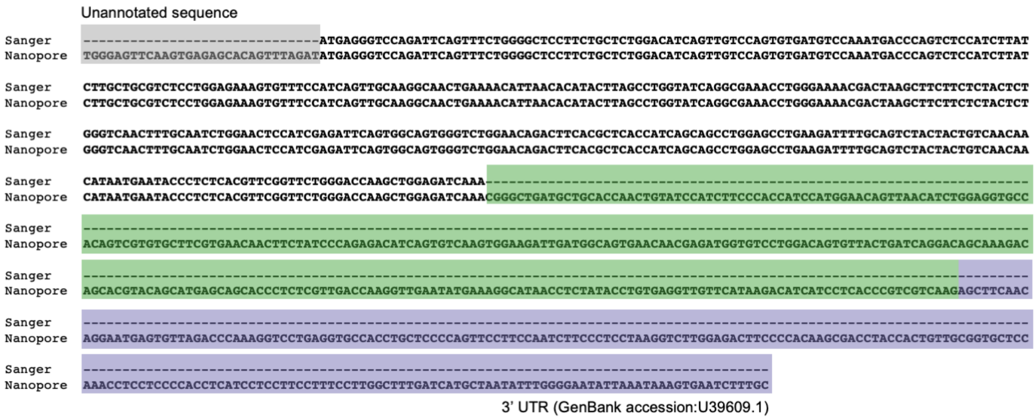


**Supplementary Figure 2.** Alignment of the light chain sequence of 7D10, as derived from Sanger (top row) and Nanopore (bottom row) sequencing methods. Additional bases present in the Nanopore sequence have been annotated with BLAST and highlighted as in Fig. 3a. The additional 5' sequence was unable to be annotated with BLAST, nor did it match any of the primer sequences used during library preparation.

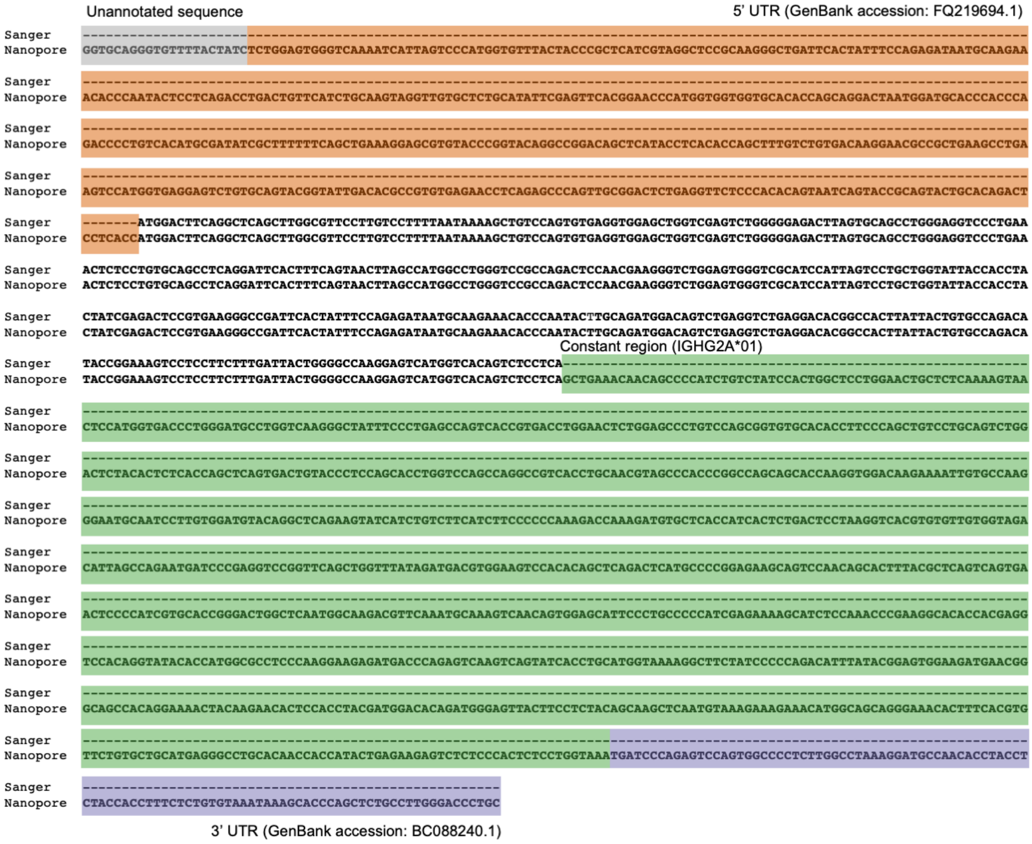


**Supplementary Figure 3.** Alignment of the heavy chain sequence of 7D10, as derived from Sanger (top row) and Nanopore (bottom row) sequencing methods. Additional bases present in the Nanopore sequence have been annotated with BLAST and highlighted as in Fig. 3a. The additional 5' sequence was unable to be annotated with BLAST, nor did it match any of the primer sequences used during library preparation.
